# leafing phenology and insect seasonality in an ever-wet tropical forest

**DOI:** 10.1101/2025.10.10.681743

**Authors:** DL Forrister, SL Tarco, DA Donoso, NC Garwood, R Valencia, PD Coley, M-J Endara

## Abstract

Seasonal patterns in tropical insect communities remain poorly understood because temperature is rarely limiting to growth or reproduction in tropical climates. In these regions, insect seasonality is often attributed to rainfall, yet distinct seasonal dynamics also occur in ever-wet forests where climatic fluctuations are minimal. Here, we examine the seasonality of abundance and diversity in lepidopteran herbivores associated with the tropical tree genus Inga in Yasuní, an aseasonal lowland rainforest in the Ecuadorian Amazon. We find that both leaf production and herbivore communities show pronounced seasonal peaks during the driest and brightest months of the year. Using structural equation modeling, we evaluate the relative roles of climate and leaf production in shaping insect seasonality and find support for both pathways. Finally, we show that communities feeding on expanding versus mature leaves differ strongly in composition, with expanding leaves supporting more specialized and phylogenetically divergent herbivore assemblages. Together, these results demonstrate that even in an ever-wet tropical forest, subtle climatic cues and host phenology interact to drive pronounced insect seasonality

## Introduction

Phenology asks a simple but fundamental question: *When do biological events such as emergence, growth, dispersal, and reproduction occur, and why at those times?* In temperate forests, these events are strongly seasonal, following annual cycles of temperature, day length (solar radiation), and precipitation. While these patterns are often taken for granted in temperate zones, the factors that shape plant and animal phenology in the tropics are far less intuitive. For example, it remains puzzling why insects in tropical regions—where cold winters are absent—maintain seasonal life cycles (Wolda, 1988). Broad-scale analyses show that activity patterns of plants and animals tend to become less seasonal with decreasing latitude and increasing rainfall (Reich, 1995). Nevertheless, studies of insect seasonality in the tropics consistently find marked fluctuations in invertebrate abundance(Kishimoto-Yamada & Itioka, 2015; Wolda, 1988).

In tropical ecosystems, insect seasonality is most often attributed to rainfall. Seasonality is especially pronounced in tropical dry forests and savannas, where insect activity typically peaks either in the wet season (Wolda, 1978, 1980, 1989; Wolda et al., 1998; Wolda & Chandler, 1996) or in the dry season (Michel & Cadet, 2009; Slaa, 2006; Wolda, 1978, 1989). By contrast, fewer studies have examined tropical wet forests, which lack a severe dry season. Results from these ecosystems reveal both seasonal (Checa et al., 2019; Devries & Walla, 2001; Grøtan et al., 2012; Wolda et al., 1998; Wolda & Chandler, 1996) and aseasonal (Novotny et al., 2002, p. 200; Wolda & Galindo, 1981) patterns of insect abundance. Given that virtually all phenological strategies occur in wet tropical forests(Kishimoto-Yamada & Itioka, 2015), a central question emerges: why do some insect species maintain a seasonal life cycle in ecosystems lacking strong abiotic constraints on growth?

Most work has focused on climatic drivers such as temperature, photoperiod, and rainfall, which can limit insect development or serve as cues for synchronized reproduction. Yet insect populations are also shaped by biotic factors, including resource availability, predation, and parasitism. Food availability is particularly critical for phytophagous insects, whose reproduction and development depend on host plants. In tropical forests, ∼80% of lifetime leaf damage occurs during leaf development, with mature leaves experiencing relatively little herbivory during their 1–3 year lifespan (Brenes-Arguedas et al., 2006; Coley & Aide, 1991; Kursar & Coley, 2003). Expanding leaves are both nitrogen-rich and less structurally defended, allowing herbivores to maximize growth and minimize exposure to predators and parasitoids (Benrey & Denno, 1997; Roslin et al., 2017). Caterpillars feeding on expanding leaves grow more than twice as fast as those restricted to mature leaves (Coley et al., 2006). Moreover, tropical herbivores are highly host-specific: most lepidopteran species feed on only 1–3 host plants (Coley et al., 2018; Endara et al., 2017; Forister et al., 2015; Forrister et al., 2019). This host specialization implies that each insect species should closely track the leafing phenology of its particular hosts.

Taken together, these findings suggest that the timing and duration of new leaf production could strongly shape insect seasonality, even in ever-wet tropical forests. Here, we examine this relationship in Yasuní, a lowland rainforest in eastern Ecuador. Yasuní lies on the equator, with nearly constant day length and temperature, and no months receiving less than 100 mm of rainfall. We test four hypotheses: (1) leaf production is seasonal despite the absence of a dry season; (2) the timing and duration of leaf production predict insect abundance and diversity; (3) herbivore communities differ between expanding and mature leaves of the same host species; and (4) herbivore species associated with expanding leaves are more specialized and display stronger seasonality.

## Materials and Methods

### Study site

All sampling was conducted at the Estación Científica Yasuní (ECY), maintained by the Pontificia Universidad Católica del Ecuador, and located within Parque Nacional Yasuní (76°23′50″ W; 0°40′27″ S). Elevation ranges from 220–260 m. The habitat is predominantly primary forest, with small clearings near the station and minor edge effects along the gravel access road. Yasuní is among the most diverse forests in the world, with 1104 tree species recorded within 25 ha (Valencia, Condit, et al., 2004; Valencia, Foster, et al., 2004). Fieldwork was conducted within the 50-ha Yasuní Forest Dynamics Plot (YFDP, established in 1995), in which all trees ≥1 cm diameter have been mapped, tagged, and identified.

### Monthly census of leaf production, herbivores, and herbivory

We monitored saplings of 15 focal *Inga* species along trails within the YFDP. *Inga* was chosen because it is one of the most abundant and species-rich genera in the plot, comprising >45 congeners and ∼6% of stems (Valencia, Condit, et al., 2004; Valencia, Foster, et al., 2004). Focal species were selected because they were sufficiently abundant to allow monitoring of 50 individuals each, ranging from moderately rare to very common. Collectively, these species represent the full range of leafing phenological strategies, from continuous to highly seasonal and synchronous.

Each individual was visited monthly from September 2018 to March 2020. The census was paused due to the global COVID-19 pandemic and then resumed for three months Aug 2021 to Oct 2021. Over times some individuals were lost or died and we replaced them when possible to achieve our target of ∼50 individuals per species. Species were distributed along trails within the plot. At each census we: (1) visually inspected all leaves for insect herbivores, (2) recorded the number of expanding leaves and new leaves produced since the previous census, and (3) marked all expanding leaves with fine metal wires, scoring herbivore damage (estimated as the percent area lost) once leaves had fully expanded at the following census. Although *Inga* hosts multiple insect guilds (Coleoptera, Orthoptera, Coreidae, Diptera, Tenthredinoidea, Phasmatodea, and Lepidoptera; (Endara et al., 2017), we focused on Lepidoptera because they are both the most abundant and the primary source of leaf damage(Barone, 1998; Janzen, 1988; Novotny et al., 2004). All feeding caterpillars were collected for DNA barcoding using cytochrome oxidase I (COI).

### Arthropod trap sampling

To sample adult Lepidoptera we used one BioQuip Universal Black Light trap near ECY and four standard Malaise traps within the YFDP. The light trap consisted of a 12W black light mounted between three clear plastic panes, positioned above a 5-gallon bucket containing 3–4 cm of 96% ethanol, and surrounded by a 0.5-inch mesh wire cage to exclude large insects. The trap was controlled by a timer cycling on for 30 min and off for 1 h, for a total of 3 h operation between 21:00–06:00. It was run for four nights per month during the new moon (<10% illumination). After each night, samples were removed and ethanol replaced.

Malaise traps were opened on the first day of each monthly census and samples collected 8 ± 1 days later, with ethanol replaced at each visit. All samples were preserved in 96% ethanol and frozen at −8 °C before transport to the University of Utah, where they were sorted and DNA-barcoded. We placed Malaise traps at four locations within the YFDP, two along the ridges and two within the valley/flood plains. At each locations we placed a canopy and a understory malaise trap.

### DNA barcoding

Morphological identification of Neotropical Lepidoptera larvae is difficult, so we used DNA barcoding for all herbivore collections. We amplified a 461 bp fragment of COI using primers BF3 (Elbrecht et al., 2017) and BR2 (Elbrecht et al., 2019) (Table S1). Caterpillar samples were sequenced individually. Malaise samples were randomly subsampled into four subsamples per trap, and light trap samples into eight subsamples. All Lepidoptera in each subsample were counted, homogenized, and processed for barcoding.

Extractions followed a direct PCR method for speed and cost-efficiency (Thongjued et al., 2019). Caterpillars were crushed in 20 µl phosphate-buffered saline (PBS) in 0.2 ml PCR tubes and incubated at 95 °C for 2 min; 1.25 µl of lysate was used as PCR template. The same procedure was used for Malaise subsamples. Light-trap samples, which contained more individuals, were extracted with EconoSpin® All-In-One Silica Membrane Mini Spin Columns. PCRs were performed using Platinum Direct PCR Universal Master Mix (ThermoFisher Scientific, USA) (Tables S2, S4).

We used a nested barcoding protocol(Kitson et al., 2019) involving two rounds of PCR with unique index combinations (Table S3), allowing pooled sequencing on an Illumina MiSeq with 2 × 300 V3 chemistry.

### Climate data

Climate data were obtained from the ECY weather station and the Tropical Ecology Assessment and Monitoring (TEAM) Network, a collaboration among Conservation International, Missouri Botanical Garden, Smithsonian Institution, and Wildlife Conservation Society, with partial funding from the Gordon and Betty Moore Foundation. Data included daily mean, minimum, and maximum air temperature, and daily rainfall (manual rain gauge). Solar radiation was not available during the study period, so we estimated photosynthetically active radiation (PAR) from MODIS product MCD18A2 V6(D. Wang & Liang, 2019).

To validate the MODIS PAR estimates, we compared them with ground-based solar irradiance at ECY for 359 overlapping observations (Jan 2018–Aug 2019). MODIS estimates were well correlated with station irradiance (adjusted R^2^ = 0.61; Fig. S1) and captured the same seasonal pattern.

## Results

### Yasuní climate

Temperature, rainfall, and solar irradiance during the study period closely tracked long-term averages for the site. Both temperature (CV = 0.03, ρ = 0.01, Rayleigh test = ns) and daylength (CV = 0.002, ρ = 0, Rayleigh test = ns) were aseasonal (Fig. 2A–B). Yasuní lies at 0.5° S and experiences nearly constant 12 h daylength (± 5 min) year-round. Photosynthetically active radiation (PAR), estimated from MODIS satellite imagery, was more seasonal (CV = 0.16, ρ = 0.07, Rayleigh test < 0.001), peaking between August and November.

Rainfall ranged from 200–400 mm per month with a unimodal peak from March to July (CV = 0.34, ρ = 0.15, Rayleigh test < 0.001). Importantly, no month received <100 mm of rain, confirming Yasuní as an “ever-wet” Amazonian forest (Lopes et al., 2016; Saleska et al., 2016; J. Wang et al., 2020). Overall, Yasuní has one of the least seasonal climatic regimes in the Neotropics. Based on temperature, rainfall, and daylength, there appear to be no strong abiotic constraints on insect growth and reproduction.

### Understory census of leaf production and insect herbivores

We marked and monitored 907 saplings from 15 focal *Inga* species (60 ± 7 individuals per species). We conducted 21 monthly censuses: 18 consecutive months from September 2018 to February 2020, followed by three additional censuses (Aug–Oct 2021) after a pandemic-related closure of Yasuní National Park. In total, we completed 15,665 sapling surveys (732 ± 41 individuals per month). Each census required ∼8 days and ∼168 person-hours.

Despite the lack of a marked dry season, leaf production was strongly seasonal, with a major flush from August–October (CV = 0.46, ρ = 0.1, Rayleigh test < 0.001) and a smaller peak in April (Fig. 1E). This pattern, with flushing during the driest and brightest months, mirrors Amazon-wide trends from satellite imagery (Lopes et al., 2016, p. 201; Saleska et al., 2016, p. 201; J. Wang et al., 2020). Most focal *Inga* species followed this general pattern (Fig. S2), though the degree of synchrony varied among species.

**Figure 1:**
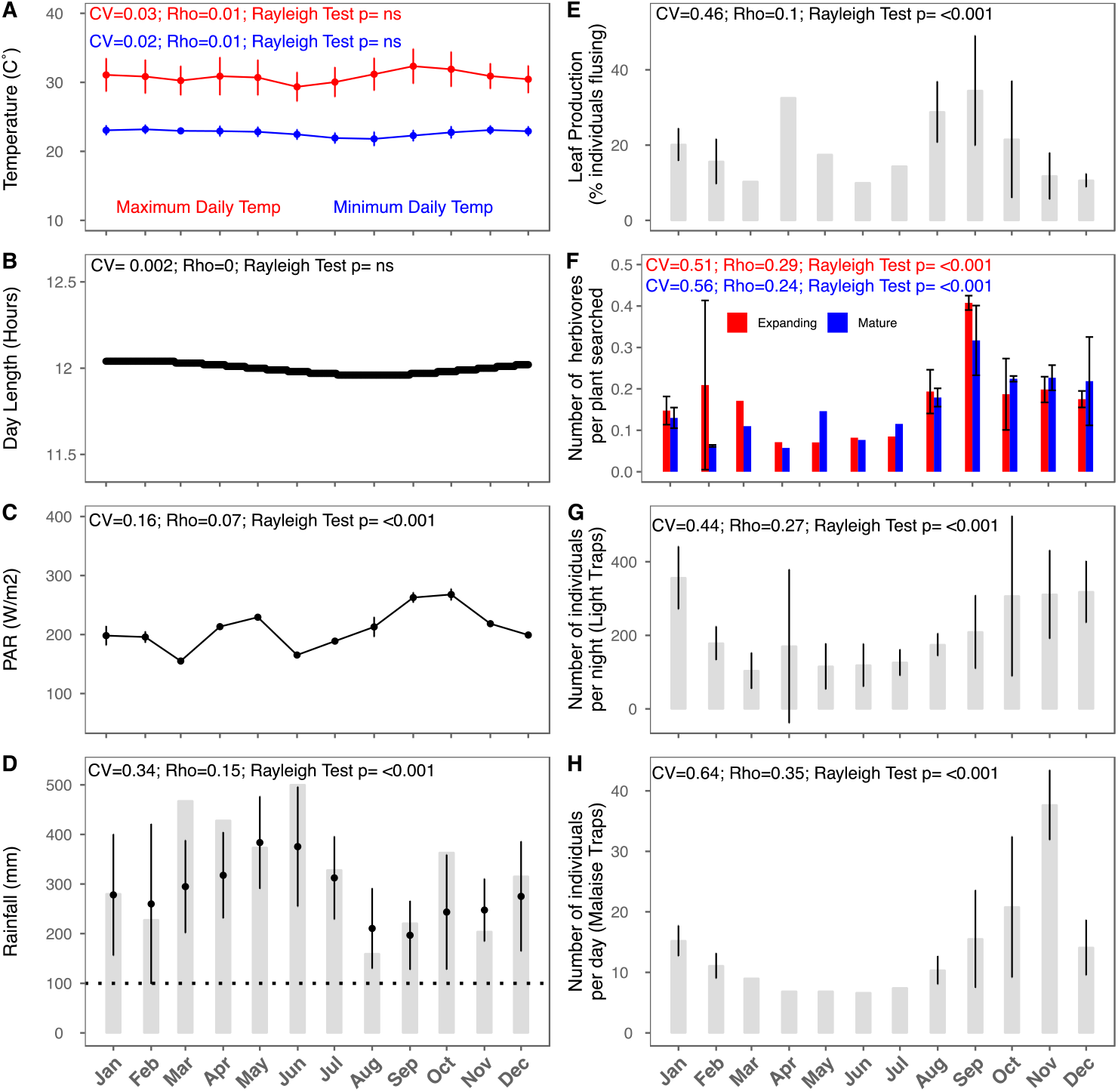
Average monthly climatic conditions, leaf production and insect abundance. A) average minimum and maximum daily temperature. B) Day length. C) Photosynthetically Active Radiation (PAR) measured from MODIS satellite imagery. D) Cumulative monthly rainfall measured daily using manual rain gage. E) Leaf production measured as the percent of individuals with expanding leaves in each monthly census. F) Number of caterpillars observed feeding during census. G) Number of individuals collected per night in light traps. H) Number of individuals per day collected in Malaise traps. Coefficient of Variation (CV), Rho’s vector, (resultant length of circular mean angle vector) and the significance of a Rayleigh test reported in each panel.

### Insect seasonality

Both caterpillar and adult Lepidoptera exhibited pronounced seasonality, with unimodal peaks in abundance from July to November (Fig. 1F–H). Across surveys we recorded 3,824 feeding caterpillars. DNA barcoding yielded COI sequences for 1,726 individuals (45% recovery), revealing 217 MOTUs across 13 Lepidopteran families.

Malaise traps (n = 4) collected 2,534 adult Lepidoptera, with monthly catches ranging from 6–41 individuals per trap-day. Light traps (12 h per month) captured far more individuals (total = 18,358; monthly range = 36–794 individuals per night). DNA metabarcoding of Malaise samples identified 575 MOTUs. All collection methods showed consistent seasonal patterns, with peaks in both abundance (Fig. 1F–H) and species diversity (Fig. 2) from September to January.

**Figure 2:**
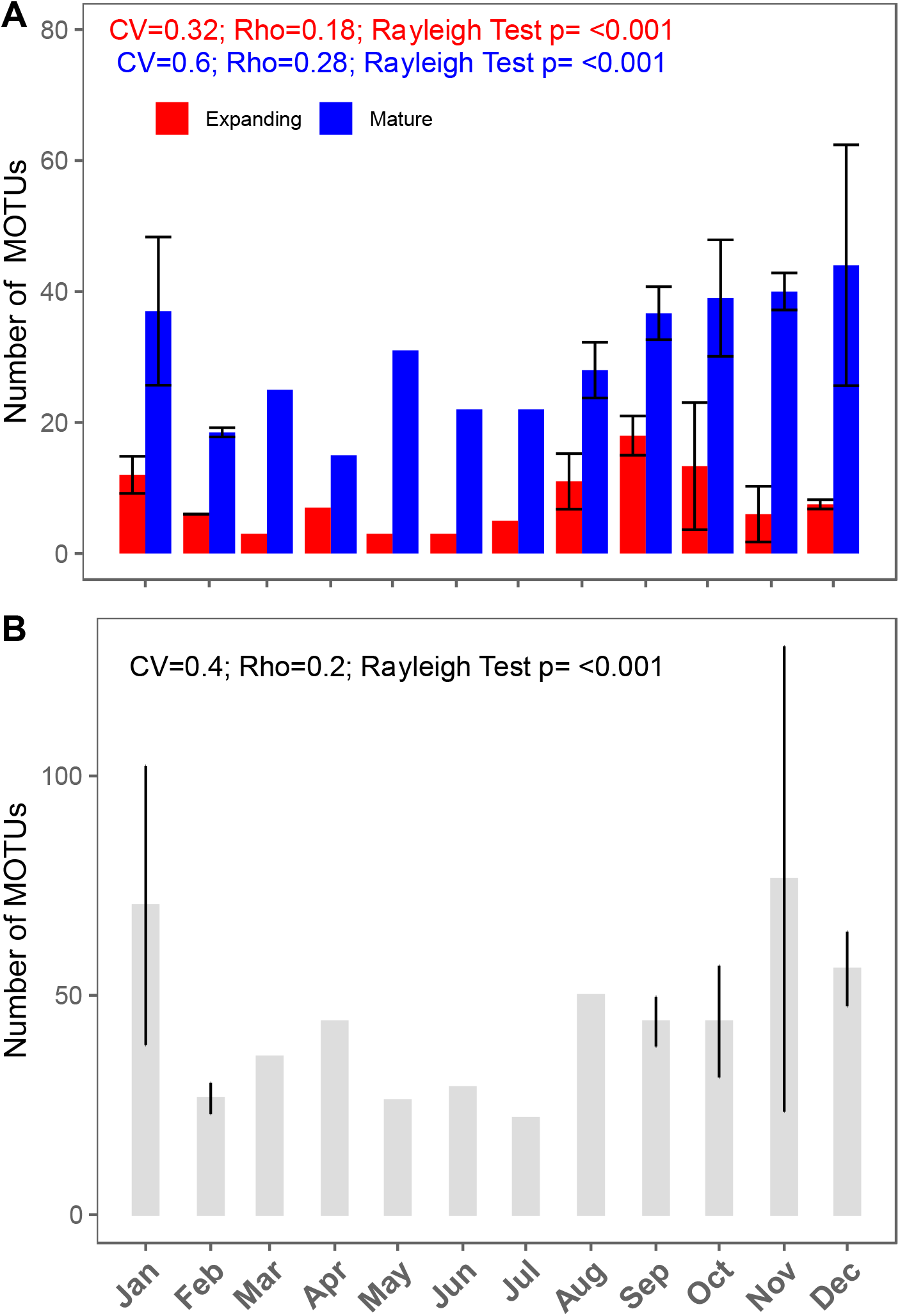
Insect diversity. A) Number of molecular operational taxonomic units (MOTUs) observed on expanding and mature leaves during each monthly census. B) Number of MOTUs observed in each month of Malaise trap sampling.

To test proximate drivers of insect seasonality, we used structural equation modeling (SEM) (Fig. 3). We first modeled climate effects (temperature, rainfall, PAR) on leaf production (% individuals flushing; GLM, binomial). PAR had a strong positive effect, while temperature had a weaker negative effect. We then modeled caterpillar abundance (GLM, Poisson) as a function of leaf production, temperature, and rainfall. Leaf production and temperature were positively correlated with caterpillar abundance, while rainfall was negatively correlated. Adult Lepidoptera abundance (GLM, Poisson) was positively correlated with both temperature and caterpillar abundance. A parallel SEM for caterpillar diversity found fewer significant effects: leaf production was primarily associated with PAR, and caterpillar diversity correlated only with adult diversity and leaf production (Fig. 3B).

**Figure 3:**
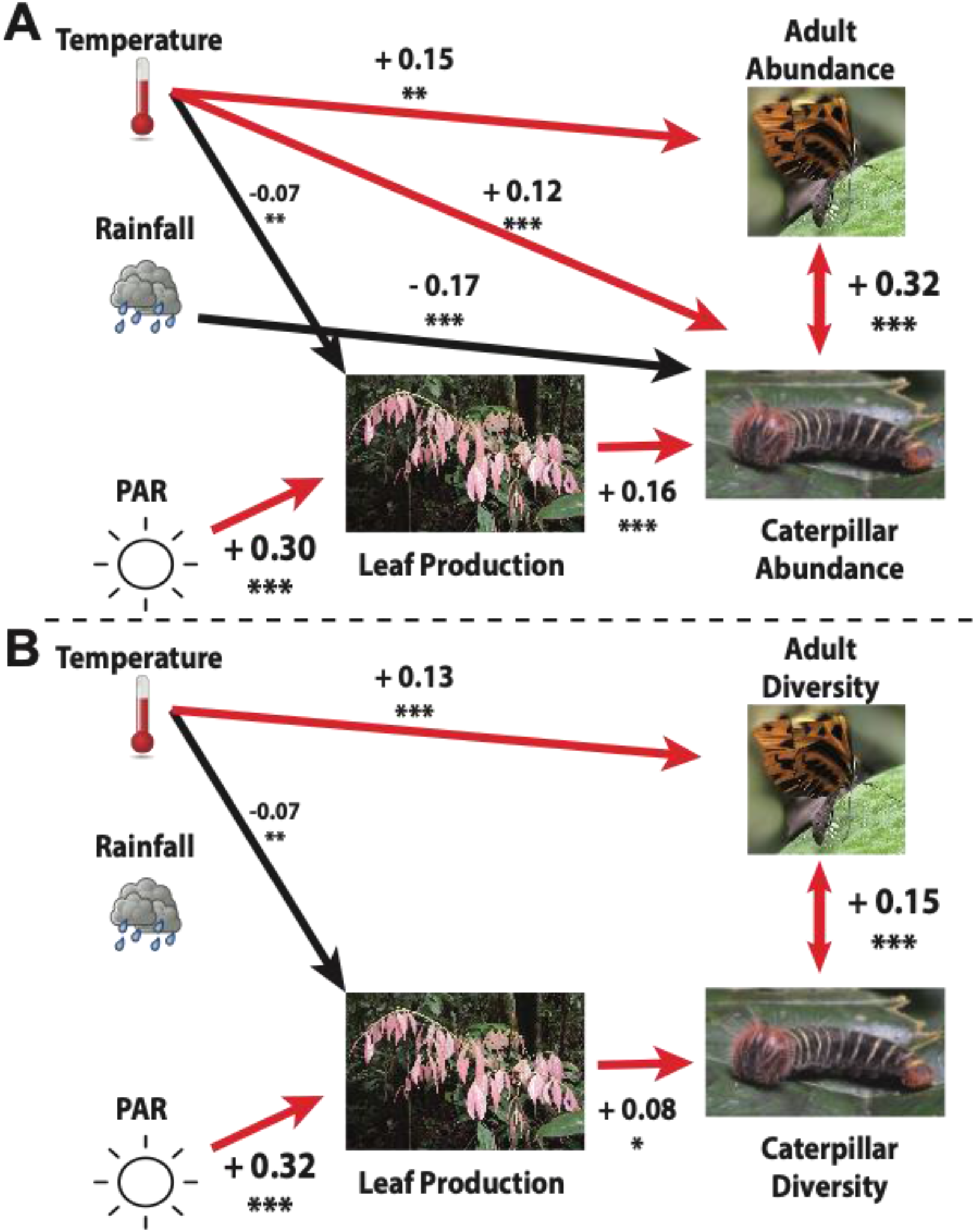
Path analysis of A) insect abundance and B) insect diversity.

### Seasonality in community composition of insect herbivores

To evaluate seasonal turnover in herbivore communities, we used principal coordinate analysis (PCoA) based on Bray–Curtis distances across the 21 censuses (Fig. 4). The first two axes explained 16.4% and 12.8% of variation, respectively. Sampling months clustered by rainfall, with the wettest months (April–July) grouping together. Months of peak leaf flushing separated along PCoA axis 1. PERMANOVA confirmed that herbivore community composition was significantly associated with rainfall (R^2^ = 0.09, p = 0.006), but not with leaf production.

**Figure 4:**
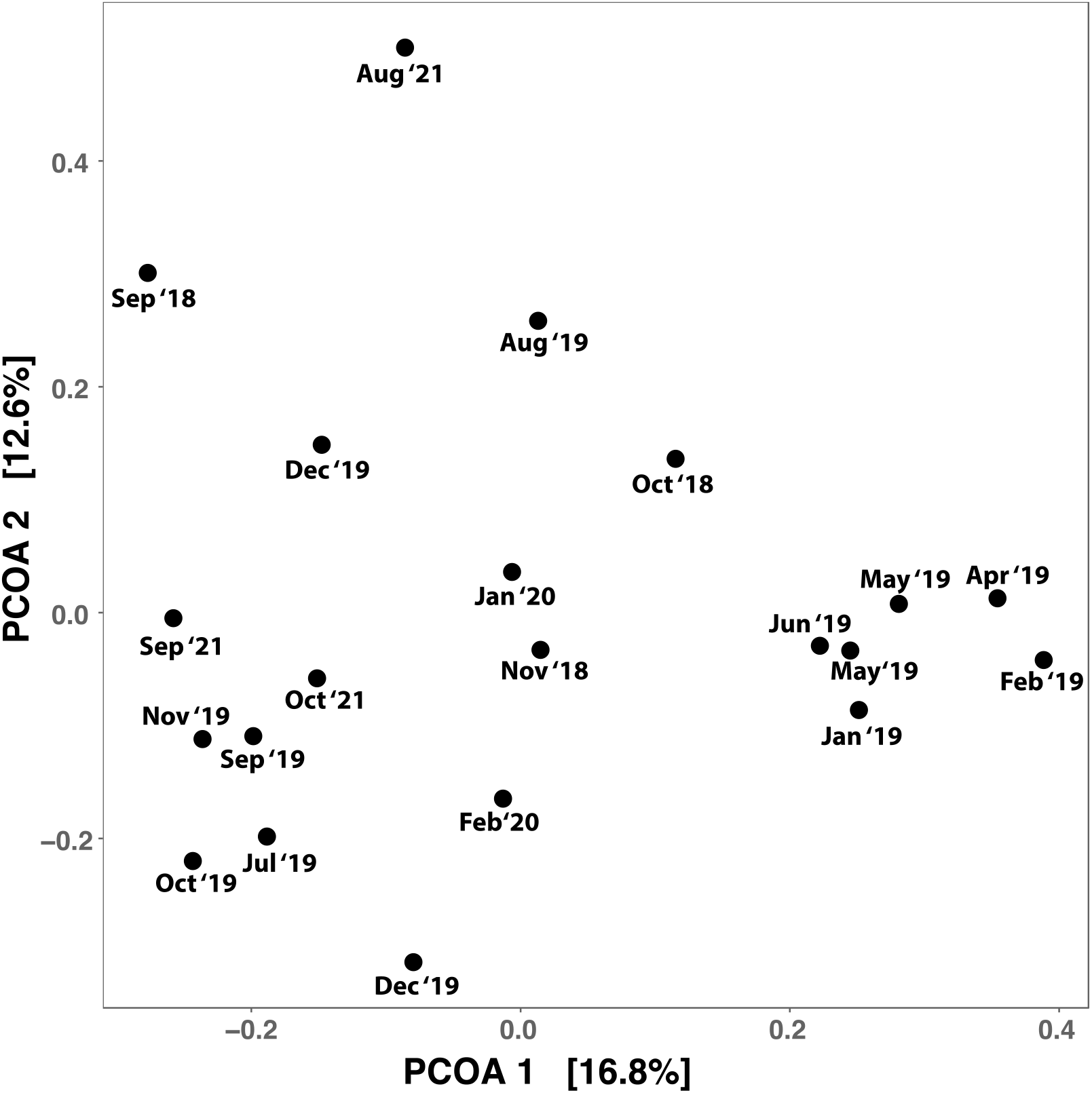
Principal coordinate analysis comparing herbivore community composition of caterpillars observed feeding on *Inga* sapling during each census.

### Herbivore communities on expanding vs. mature leaves

Tropical insect herbivores are highly specialized, typically restricted to 1–3 host species, often keyed to specific plant defenses(Endara et al., 2018). While specialization across host species is well documented, the degree of specialization on different leaf stages is less clear. Expanding and mature leaves differ sharply in chemistry (Wiggins et al., 2016), suggesting they may support distinct herbivore communities.

Constrained ordination, controlling for host species, confirmed strong compositional differences between expanding and mature leaves (PERMANOVA: p = 0.002, R^2^ = 0.11; Fig. 5). Mature leaves harbored more individuals and more species overall (Fig. 6). However, herbivores associated with expanding leaves were more specialized: they used fewer host species and displayed greater divergence among hosts (Fig. 7).

**Figure 5:**
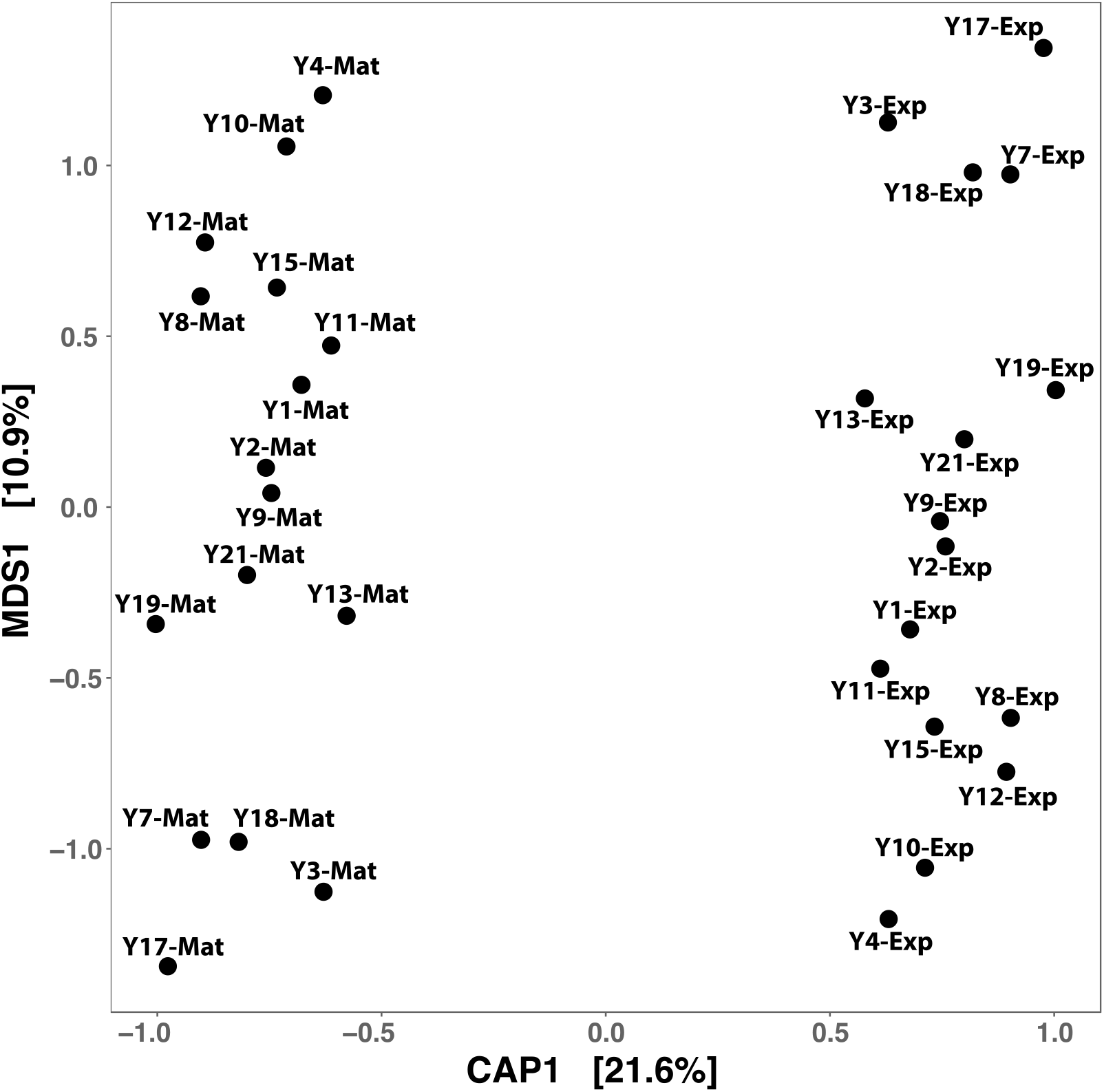
Constrained Analysis of Principal coordinates comparing the composition of herbivore communities between expanding (Exp) and mature (Mat) leaves, conditioned on *Inga* host species. Numbers indicate different *Inga* species.

**Figure 6:**
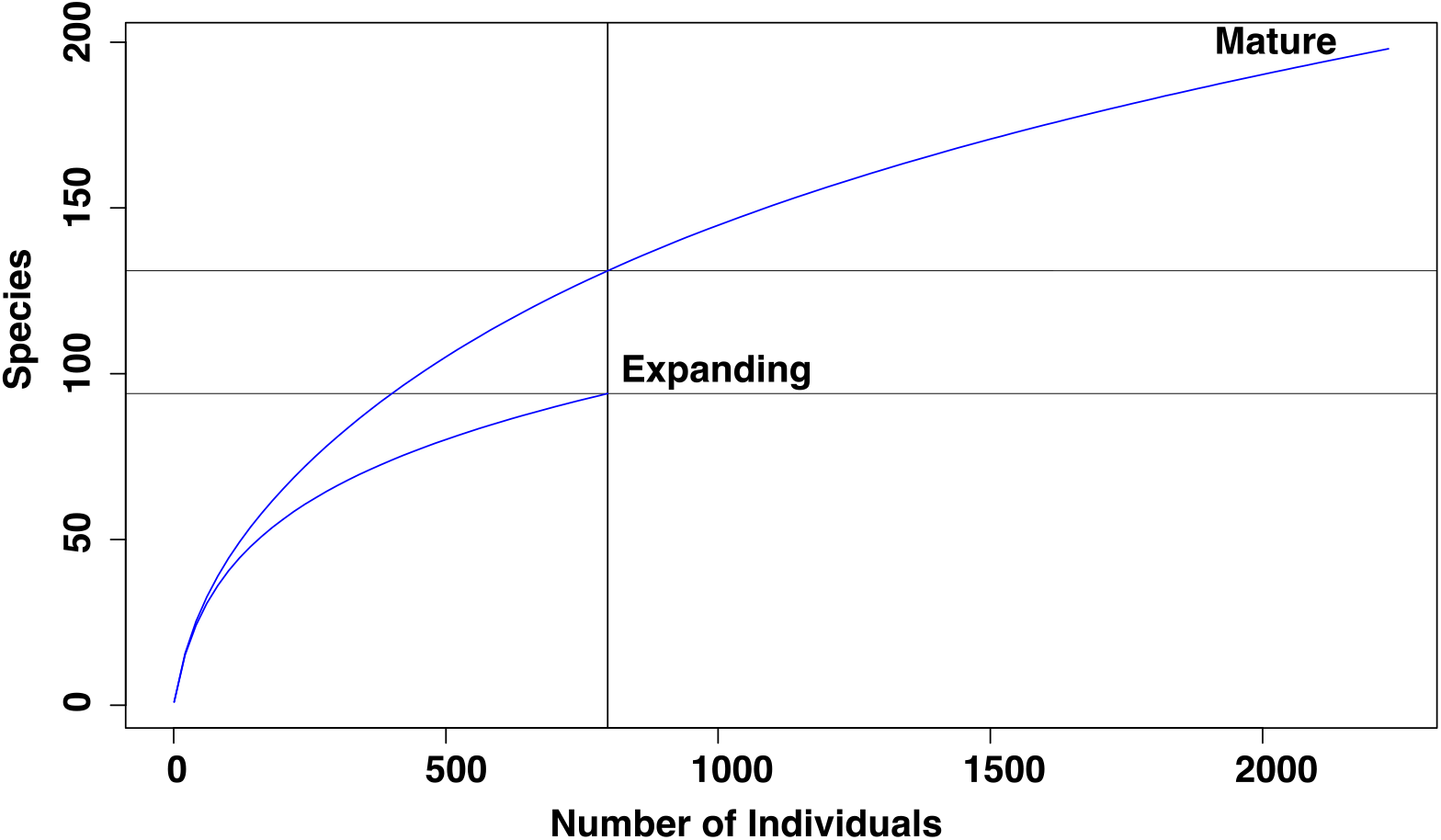
Rarefaction of number of herbivore MOTUs collected on expanding and mature leaves during all 21 census months.

**Figure 7:**
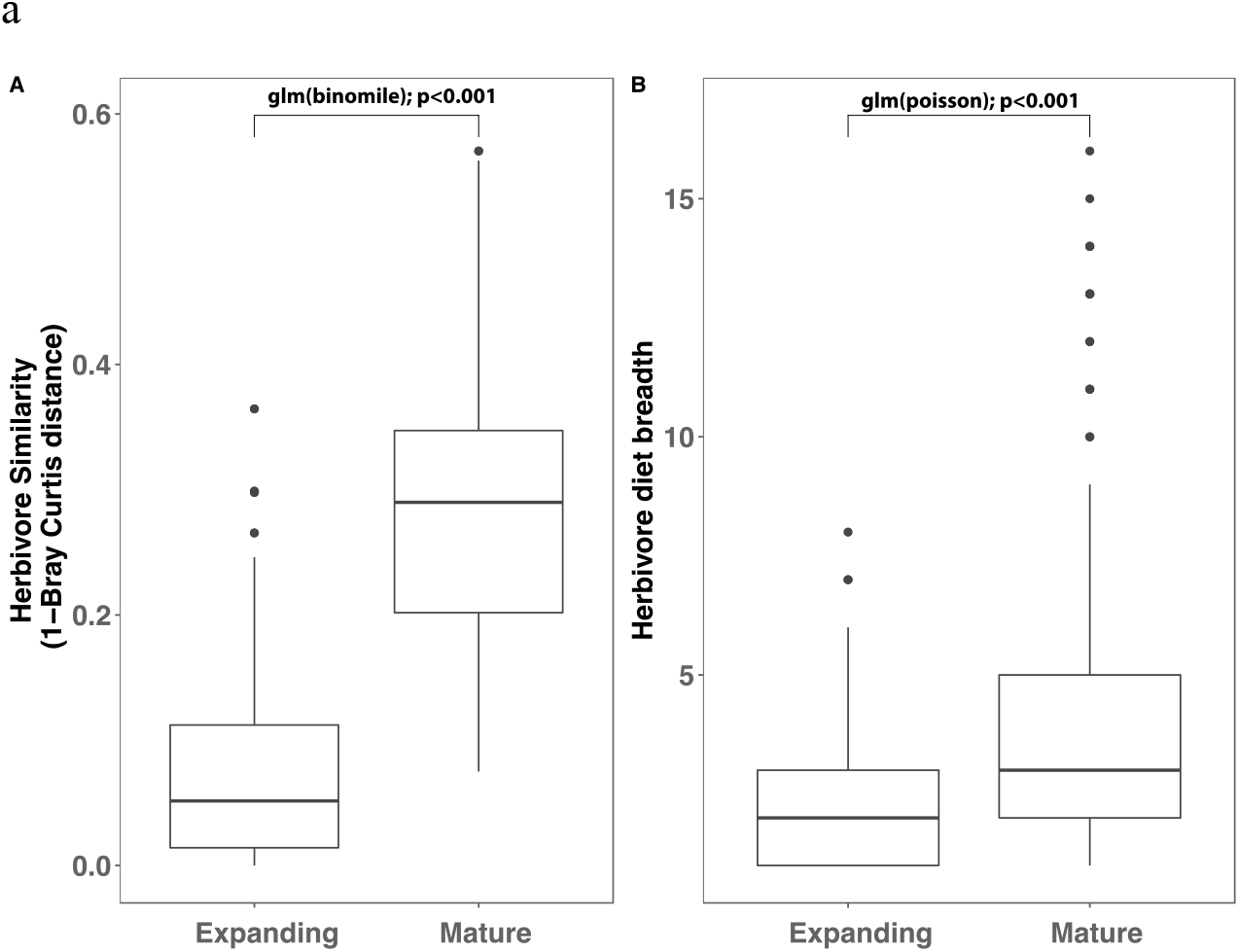
Comparison of expanding and mature leaf herbivore communities in terms of A) similarity (1-Bray Curtis dissimilarity) between *Inga* hosts and B) herbivore diet breadth (number of host species).

## Discussion

Yasuní, Ecuador, lies in the “Core Amazon,” one of the wettest regions of the basin with no severe dry season (Killeen & Solorzano, 2008). Based on global syntheses, insect seasonality is more prevalent in tropical dry forests than in wet forests (Kishimoto-Yamada & Itioka, 2015). Our findings challenge this expectation: we observed pronounced seasonal peaks in the abundance and diversity of lepidopteran herbivores in Yasuní, one of the least seasonal regions of the Neotropics. Although temperature and rainfall do not impose strong physiological constraints on insect growth, both rainfall and leaf production showed marked seasonal patterns, and these factors helped explain insect seasonality at this site. In contrast, many Southeast Asian forests experience weaker seasonality in both climate and phenology, in part because rainfall is higher and less predictable, lacking a consistent annual peak (Kishimoto-Yamada & Itioka, 2015). The year-to-year consistency of rainfall and solar irradiance in Yasuní may provide a reliable cue that plants and insects can track. Our results, showing peak lepidopteran diversity and abundance during the brightest and driest months of the year, are consistent with a butterfly-focused study from the same site (Checa et al., 2009).

Most research on insect seasonality in the tropics has focused on adults, sampled via traps (Kishimoto-Yamada & Itioka, 2015; Wolda, 1988). By focusing on larvae, we were able to explicitly link insect abundance and diversity to leafing phenology. Only a handful of previous studies have made this connection (Ichie et al., 2004; Itioka & Yamauti, 2004; Wolda, 1978). We found that leaf production was a strong predictor of both insect abundance and diversity. Thus, even in an ever-wet Amazonian forest, phenological synchrony between expanding leaves and herbivore activity appears to be a key driver of insect seasonality. An exception was the April leaf flush, which was not accompanied by a peak in herbivores. This may reflect differences between canopy and understory phenology, as canopy trees often show a single annual flush, while understory individuals may flush multiple times per year (Barone, 1996).

Mature-leaf herbivores also peaked during the brightest and driest months. Although mature leaves are available year-round, higher adult activity and oviposition during this period may generate seasonal pulses in mature-leaf herbivores. We also observed a 1–2 month lag between peak leaf flushing and peak mature-leaf herbivore abundance, consistent with the higher nutritional quality of recently expanded leaves compared with older foliage (Coley & Kursar, 1996; Kitajima et al., 1997).

These results fit into a growing consensus that leaf flushing is seasonal across the Amazon basin, driving a basin-wide “green-up” at the onset of the dry season(J. Wang et al., 2020; Wu et al., 2016). Our understory data from Yasuní match this regional pattern. Moreover, we show that seasonal variation in leaf production cascades upwards to structure herbivore communities, reinforcing the importance of plant phenology as a driver of tropical forest ecology.

Diversity patterns mirrored abundance patterns, with both caterpillars and adults peaking during the driest, brightest months. The flush of new leaves provides an ephemeral niche, supporting a distinctive community of herbivores. However, diversity on expanding leaves was lower than on mature leaves after controlling for individual abundance. Expanding leaves are more chemically defended (Coley & Kursar, 1996; Wiggins et al., 2016) and transient, supporting a smaller guild of more specialized herbivores. Indeed, we found that herbivores on expanding leaves were more host-restricted and more divergent among host species than those on mature leaves.

These findings underscore the dual importance of host species identity and leaf developmental stage in shaping herbivore specialization. Previous studies have shown that divergence in host defenses drives host specialization(Endara et al., 2017, 2018; Forrister et al., 2019), and our results extend this pattern to intra-individual variation in defense across leaf ontogeny. In tropical forests, high predation and parasitism(Coley et al., 2018; Roslin et al., 2017) likely interact with divergent defense traits to maintain the extreme host specificity of lepidopteran herbivores. In turn, this specialization causes herbivore species to track host phenology closely, giving rise to pronounced insect seasonality even in one of the wettest and least seasonal environments on Earth.

## Conclusion

In this study we demonstrated that both leaf production and insect communities are seasonal in Yasuní, Ecuador, one of the wettest and climatically aseasonal forests in the Neotropics. Synchronous and seasonal leaf production may be selected for because mast leafing may satiate herbivores resulting in overall lower damage(Aide, 1988). Leaf production at the start of the dry season in light limited ever-wet forests may also allow trees to maximize their photosynthetic output during the brightest months of the year (Chavana-Bryant et al., 2017; Lopes et al., 2016; Wu et al., 2016).We found that leaf production was an important predictor of insect seasonality indicating that plant defense and leaf production influence higher trophic levels. In general we know remarkably little about insect seasonality, especially in tropical wet forests(Kishimoto-Yamada & Itioka, 2015). More studies are needed for more insect groups given increasing concerns about widespread decline in insect populations (van der Sluijs, 2020). Our observation that insect seasonal dynamics are tied with plant phenology emphasizes the potential for phenological mismatch to exacerbate the effects of climate change on the health of insect populations(Renner & Zohner, 2018).

## Data Availability Statement

All data and R scripts are available upon reasonable request and will be deposited to Dryad/GitHub upon journal submission. Sequences are available in BOLD and NCBI.

## Acknowledgments

This was facilitated by the Ministerio del Ambiente del Ecuador who provided all necessary permits. Thanks to all the undergraduate assistants who worked on this project: L. Saige Alloway, Adolfo Chamba, Seung Joon Lee, Jeneth Renteria, Dayana Saraguro, Graham Taggart, Stefany Tarco, Karrin Tennent, Linda Zhao. This work was supported by the following grants: NSF GRFP, GRIP and GROW, U of U GCSC and CLAS Field Research Grant, Lewis and Clark Fund for Exploration and Field Research, ForestGEO to DLF and NSF and Smithsonian to PDC.

## Notes

### Competing Interest Statement

The authors have declared no competing interest.

