## Supplemental Data for "leafing phenology and insect seasonality in an ever-wet tropical forest"

### Supplementary materials

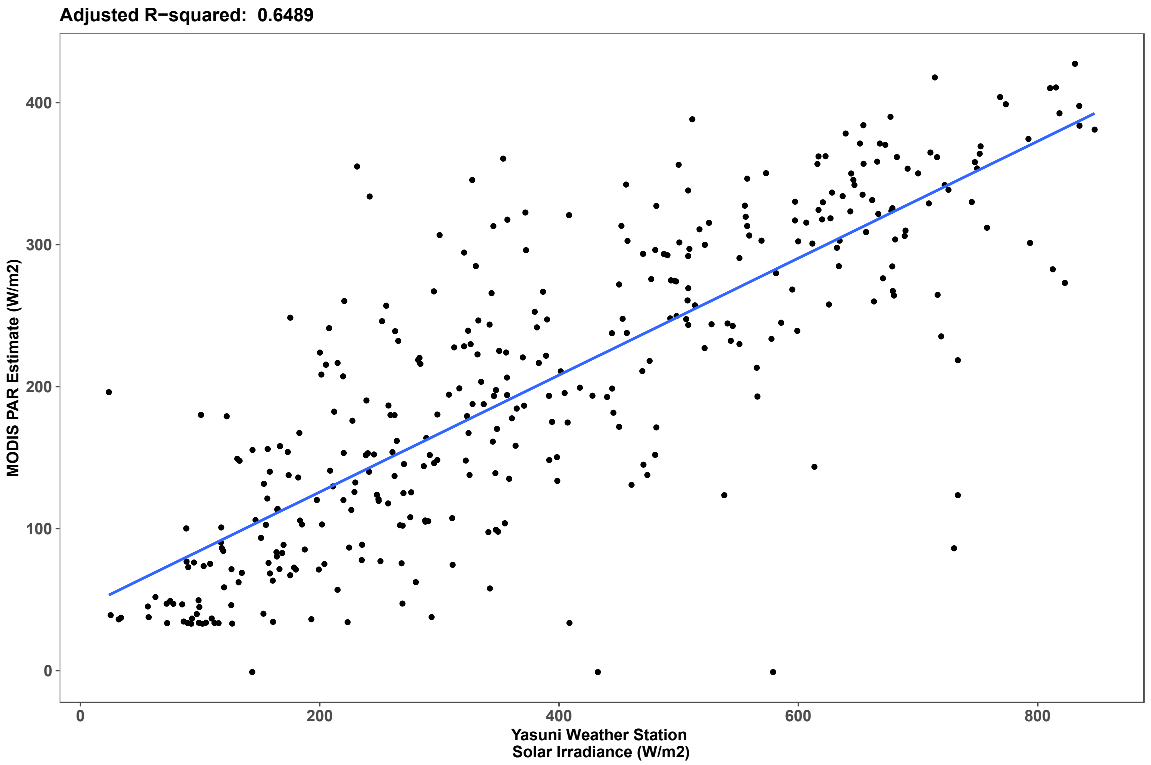

Figure S1: Correlation between Solar Irradiance measured in situ at the Estación Científica Yasuní’s weather station and photosynthetically active radiation (PAR) measured via MODIS satellite imagery.

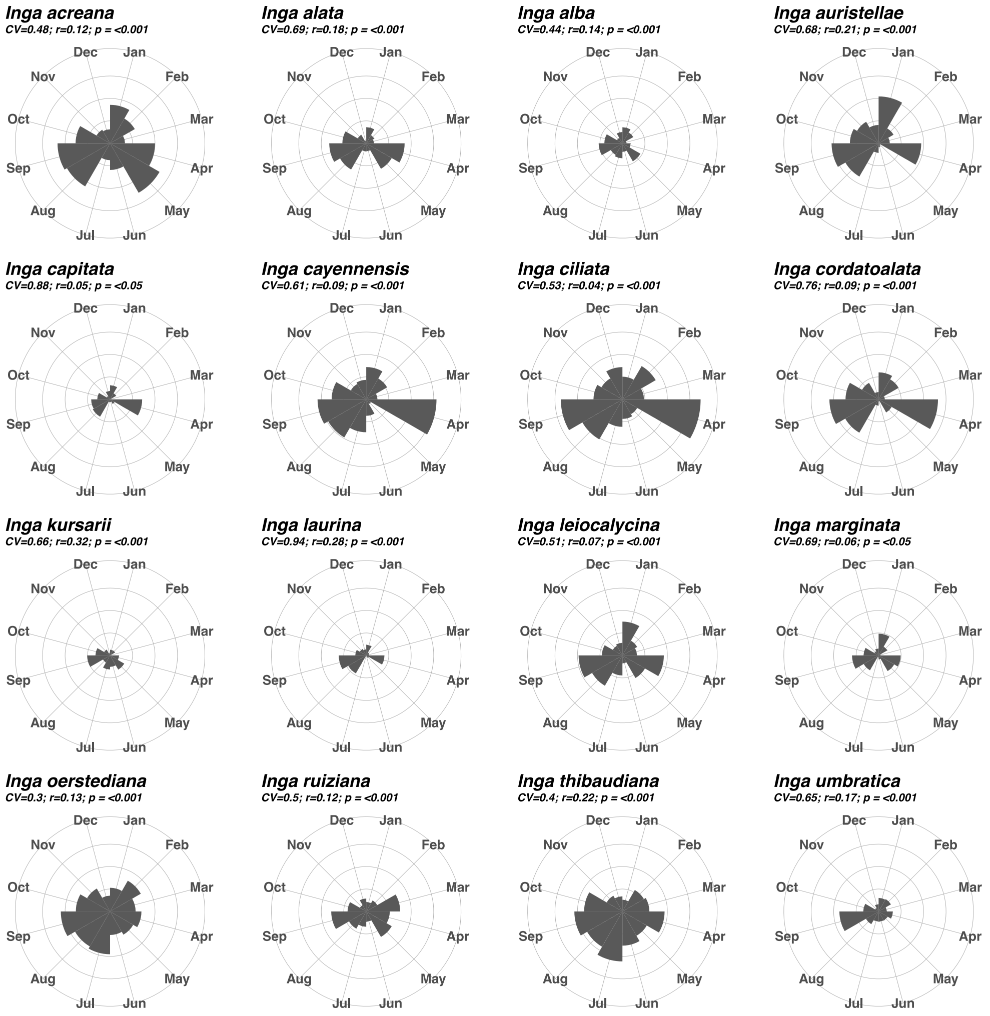

Figure S2: Leaf production by focal species, measured as the percent of individuals flushing each month.

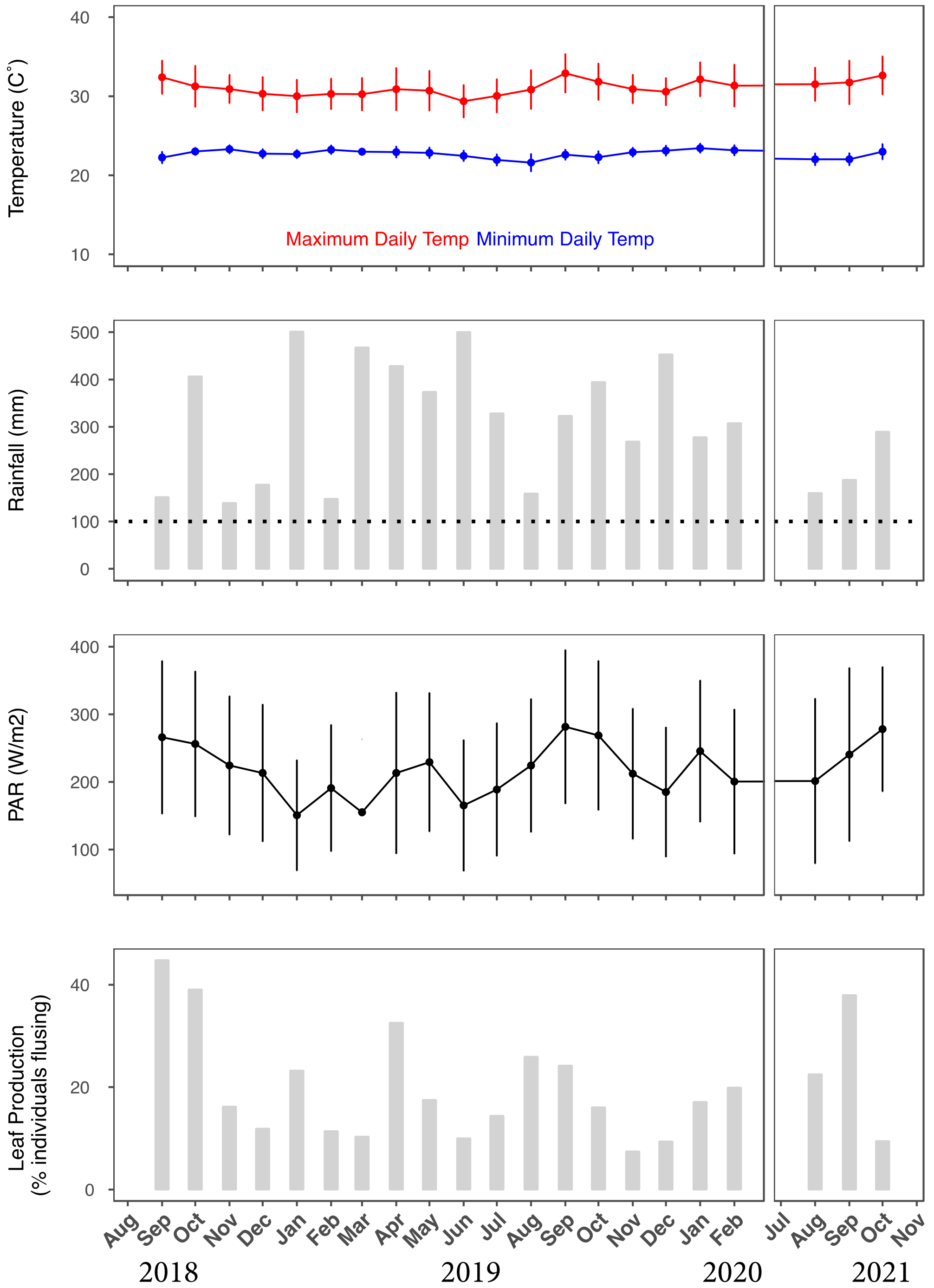

Figure S3: Climatic conditions and leaf production for each of the 21 monthly censuses conducted in this study.

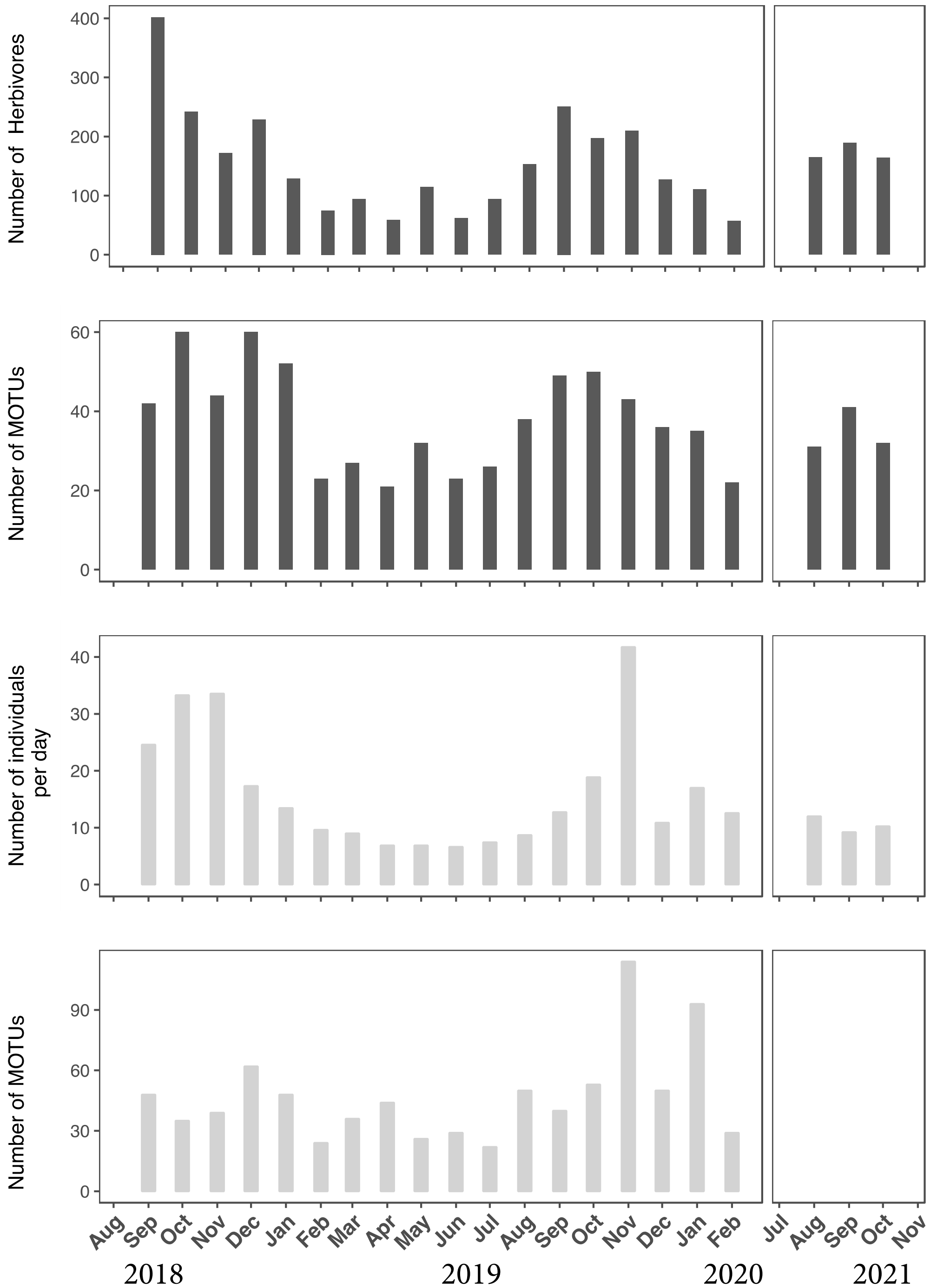

Figure S4: Abundance and diversity of insects observed during each of the 21 monthly censuses conducted in this study. A) Caterpillar abundance collected on focal saplings. B) Number of MOTUs observed in collections of caterpillars. C) Number of individuals observed in Malaise trap samples. D) number of MOTUs observed in Malaise trap samples.

**Molecular methods for DNA barcoding Cytochrome *c* oxidase subunit I (COI):**

Primers used amplify the COI barcoding region, using either BF3/BR2 (Elbrecht & Leese, 2017)). Illumina Nextera overhang adapters were added to each primer (FWD 5'-TCGTCGGCAGCGTCAGATGTGTATAAGAGACAG-3' and REV 5'-GTCTCGTGGGCTCGGAGATGTGTATAAGAGACAG-3')  In order to multiplex up to 36 plates on a single run we added unique identifiers to each primer and included variable length spaces to primers to increase base call diversity (Kitson et al., 2019).

Primers Used:

| ID | Name | Variable Spacer | Index | Sequence (5' to 3') |
|---|---|---|---|---|
| **F1_1** | BF3_0sp | | CCHGAYATR | TCGTCGGCAGCGTCAGATGTGTATAAGAGACAGCCHGAYATRGCHTTYCCHCG |
| **F1_2** | BF3_1sp | A | ACCHGAYAT | TCGTCGGCAGCGTCAGATGTGTATAAGAGACAGACCHGAYATRGCHTTYCCHCG |
| **F1_3** | BF3_2sp | CT | CTCCHGAYA | TCGTCGGCAGCGTCAGATGTGTATAAGAGACAGCTCCHGAYATRGCHTTYCCHCG |
| **F1_4** | BF3_3sp | TAG | TAGCCHGAY | TCGTCGGCAGCGTCAGATGTGTATAAGAGACAGTAGCCHGAYATRGCHTTYCCHCG |
| **F1_5** | BF3_4sp | GGTG | GGTGCCHGA | TCGTCGGCAGCGTCAGATGTGTATAAGAGACAGGGTGCCHGAYATRGCHTTYCCHCG |
| **F1_6** | BF3_5sp | GCTGC | GCTGCCCHG | TCGTCGGCAGCGTCAGATGTGTATAAGAGACAGGCTGCCCHGAYATRGCHTTYCCHCG |
| **R1_1** | BR2_0sp | | TCDGGRTGN | GTCTCGTGGGCTCGGAGATGTGTATAAGAGACAGTCDGGRTGNCCRAARAAYCA |
| **R1_2** | BR2_1sp | C | CTCDGGRTG | GTCTCGTGGGCTCGGAGATGTGTATAAGAGACAGCTCDGGRTGNCCRAARAAYCA |
| **R1_3** | BR2_2sp | AG | AGTCDGGRT | GTCTCGTGGGCTCGGAGATGTGTATAAGAGACAGAGTCDGGRTGNCCRAARAAYCA |
| **R1_4** | BR2_3sp | ATG | ATGTCDGGR | GTCTCGTGGGCTCGGAGATGTGTATAAGAGACAGATGTCDGGRTGNCCRAARAAYCA |
| **R1_5** | BR2_4sp | GAAG | GAAGTCDGG | GTCTCGTGGGCTCGGAGATGTGTATAAGAGACAGGAAGTCDGGRTGNCCRAARAAYCA |
| **R1_6** | BR2_5sp | TTGCA | TTGCATCDG | GTCTCGTGGGCTCGGAGATGTGTATAAGAGACAGTTGCATCDGGRTGNCCRAARAAYCA |

Table S1: Primers used in PCR round 1.

Table S2

| **PCR mix:** | |
|---|---|
| **Reagents** | **Volume (µl)** |
| MilliQ water | 3.8 |
| Direct PCR MM | 5 |
| primer BF3 (10µM) | 0.2 |
| primer BR2 (10µM) | 0.2 |
| DNA template | 1.25 |
| **TOTAL** | **10.45** |
| **PCR conditions:** | |
| Step 1: 94°C | 2 minutes |
| Step 2: 94°C | 15 seconds |
| Step 3: 55.2°C | 15 seconds |
| Step 4: 68°C | 22 seconds |
| Go to step 2, repeat 15x | |
| Step 5: 68°C | 6 minutes |
| Step 6: 10°C | hold |
| END | |

Table S2: PCR reagents and conditions for PCR round 1

| ID | Index | Sequence (5' to 3') |
|---|---|---|
| **F2_1** | AACTGTCC | AATGATACGGCGACCACCGAGATCTACACaactgtccTCGTCGGCAGCGTC |
| **F2_3** | ACTACGAC | AATGATACGGCGACCACCGAGATCTACACactacgacTCGTCGGCAGCGTC |
| **F2_7** | CACTAGGA | AATGATACGGCGACCACCGAGATCTACACcactaggaTTCGTCGGCAGCGTC |
| **F2_10** | CCTGTATC | AATGATACGGCGACCACCGAGATCTACACcctgtatcTTCGTCGGCAGCGTC |
| **F2_13** | CTTCAACC | AATGATACGGCGACCACCGAGATCTACACcttcaaccTTCGTCGGCAGCGTC |
| **F2_16** | GCATACGT | AATGATACGGCGACCACCGAGATCTACACgcatacgtTTCGTCGGCAGCGTC |
| **F2_19** | GTAACGGA | AATGATACGGCGACCACCGAGATCTACACgtaacggaTTCGTCGGCAGCGTC |
| **F2_22** | TCGCTAAG | AATGATACGGCGACCACCGAGATCTACACtcgctaagTTCGTCGGCAGCGTC |
| **R2_1** | AACTGTCC | CAAGCAGAAGACGGCATACGAGATggacagttGTCTCGTGGGCTCGG |
| **R2_3** | ACTACGAC | CAAGCAGAAGACGGCATACGAGATgtcgtagtGTCTCGTGGGCTCGG |
| **R2_5** | ATTCGCGA | CAAGCAGAAGACGGCATACGAGATtcgcgaatGTCTCGTGGGCTCGG |
| **R2_7** | CACTAGGA | CAAGCAGAAGACGGCATACGAGATtcctagtgGTCTCGTGGGCTCGG |
| **R2_9** | CCAACCAA | CAAGCAGAAGACGGCATACGAGATttggttggGTCTCGTGGGCTCGG |
| **R2_11** | CGTTCAAC | CAAGCAGAAGACGGCATACGAGATgttgaacgGTCTCGTGGGCTCGG |
| **R2_13** | CTTCAACC | CAAGCAGAAGACGGCATACGAGATggttgaagGTCTCGTGGGCTCGG |
| **R2_15** | GATAGTGG | CAAGCAGAAGACGGCATACGAGATccactatcGTCTCGTGGGCTCGG |
| **R2_17** | GCGTTAGA | CAAGCAGAAGACGGCATACGAGATtctaacgcGTCTCGTGGGCTCGG |
| **R2_19** | GTAACGGA | CAAGCAGAAGACGGCATACGAGATtccgttacGTCTCGTGGGCTCGG |
| **R2_21** | TCGACACT | CAAGCAGAAGACGGCATACGAGATagtgtcgaGTCTCGTGGGCTCGG |
| **R2_23** | TGCTTCCT | CAAGCAGAAGACGGCATACGAGATaggaagcaGTCTCGTGGGCTCGG |

Table S3: Primers used for PCR round 2, based on Illumina Nextera adapters.

| **PCR mix:** | |
|---|---|
| **Reagents** | **Volume (µl)** |
| MilliQ water | 3.8 |
| Direct PCR MM | 5 |
| primer i5-adapert (10µM) | 0.4 |
| primer i7-adapert (10µM) | 0.2 |
| DNA template | 1.2 |
| **TOTAL** | **10.45** |
| **PCR conditions:** | |
| Step 1: 94°C | 2 minutes |
| Step 2: 94°C | 15 seconds |
| Step 3: 60°C | 15 seconds |
| Step 4: 68°C | 22 seconds |
| Go to step 2, repeat 20x | |
| Step 5: 68°C | 5 minutes |
| Step 6: 10°C | hold |
| END | |

Table S4: PCR reagents and conditions for PCR round 1
